# Activation of a STAT3/LATS1 Signaling Axis by Folate Receptor α Enables Breast Cancer Cells to Resist Physiological Ferroptotic Stress

**DOI:** 10.64898/2026.09.17.751474

**Authors:** Prajakta Prasad Ambegaokar, Hira Lal Goel, Emmet R. Karner, Boris S. Dimitrov, Christi A. Silva, Arthur M. Mercurio

## Abstract

Understanding mechanisms that enable cancer cells to evade ferroptotic stress such as that imposed by detachment from extracellular matrix (ECM) is a significant problem that has important ramifications for tumor biology and therapy. We addressed this problem initially by analyzing single cell RNA-seq (scRNA-seq) data obtained from breast tumor organoids that had been treated with the ferroptosis inducer IKE. This bioinformatic analysis revealed that expression of folate receptor α (FRα) is increased in ferroptosis-resistant sub-populations. Subsequent experiments established a causal role for FRα in mediating resistance to ferroptosis triggered by ECM detachment, as well as by IKE treatment. The ability of FRα to resist ferroptosis is dependent on the non-canonical activation of STAT3, which has a key role in ferroptosis resistance. Our experimental data revealed that under ECM-detached conditions, ferroptosis resistant cells have increased *LATS1*, a core kinase in the Hippo pathway. We established that FRα regulates *LATS1* expression and identified a novel signaling axis that involves the regulation of *LATS1* transcription by STAT3 that results in YAP inhibition and the consequent repression of acyl-CoA synthetase long-chain family member 4 (ACSL4), a lipid-modifying enzyme that is essential for ferroptosis. Together, these results highlight an unexpected role for FRα in resisting the ferroptotic stress caused by ECM detachment and IKE treatment that is associated with its non-canonical signaling functions.

## Introduction

Dynamic changes in the microenvironment that carcinoma cells encounter during their progression can trigger ferroptosis, a regulated form of cell death characterized by the iron-dependent lipid peroxidation of cell membranes[1]. This situation is exacerbated during metastasis because tumor cells in circulation are exposed to conditions of matrix-detachment, oxidative stress and high iron concentration[2, 3]. The fact that many tumor cells can survive such conditions indicates that they have acquired mechanisms that enable them to resist ferroptotic stimuli. Understanding the nature of these resistance mechanisms is a timely and significant problem in cancer biology that has ramifications for therapy. Strategies that disable these mechanisms have the potential to improve cancer therapy, especially of metastatic disease. The metastatic journey involves processes such as detachment from the extracellular matrix (ECM) that can increase ROS and mechanisms that need to be acquired by circulating tumor cells (CTCs) to buffer ROS to enable metastatic cells to survive under these conditions [4, 5]. These observations are consistent with recent reports that tumor cells must acquire mechanisms to evade ferroptosis to metastasize efficiently [6, 7]. For these reasons, investigating how tumor cells survive ECM detachment is a powerful experimental approach to gain insight into novel mechanisms of ferroptosis resistance that are relevant to the metastatic process.

Our goal in this study was to identify specific cell surface proteins expressed on breast cancer cells that enable them to resist ferroptotic stress, especially the stress induced by ECM detachment. Initially, we used an unbiased approach that involved our analysis of single-cell RNA sequencing data (scRNA-seq) that we obtained from an organoid derived from a breast cancer patient that was exposed to a ferroptosis-inducing drug. Previously, we reported that this organoid was heterogeneous and contained populations of tumor cells that were ferroptosis resistant in comparison to the other tumor cell populations [8]. Analysis of these scRNA-seq data for cell surface proteins that are expressed on the ferroptosis resistant populations revealed folate receptor α (FRα) as a top hit, which captured our attention for several reasons. We observed that FRα expression is induced upon ECM detachment. This receptor is also of interest because it is expressed in breast cancers, especially more aggressive subtypes such as triple-negative breast cancer (TNBC) that have a poor prognosis [9–12], and it has been identified as a therapeutic target and several drugs and antibody drug conjugates are being evaluated for their efficacy, especially for ovarian cancer[13]. Its potential role in ferroptosis resistance, however, has not been investigated. Here, we provide evidence that FRα has a causal role in enabling breast cancer cells to resist the ferroptotic stress of ECM detachment, as well as IKE treatment. We describe a mechanism involving its non-canonical activation of STAT3 that regulates Hippo signaling and culminates in the repression of acyl-CoA synthetase long-chain family member 4 (ACSL4), a metabolic enzyme that is an essential component of ferroptosis because it regulates the lipid composition of membranes.

## Results

### Ferroptosis resistant breast cancer cells express elevated levels of folate receptor α (FRα)

To investigate how tumor heterogeneity influences ferroptosis sensitivity, we analyzed data from a previous study from our lab that used single-cell RNA sequencing (scRNA-seq) to identify specific sub-populations in an organoid derived from a human breast tumor that were resistant to the ferroptosis inducer, imidazole ketone erastin (IKE) [8]. Specifically, we sought to identify cell surface proteins that are associated with ferroptosis resistance by performing a differential gene expression analysis between IKE “non-responder” and “responder” cell populations. This analysis revealed a significant upregulation in the expression of the mRNA that encodes *FOLR1* (FRα) expression in non-responder or ferroptosis-resistant populations (**Fig. 1A**). We validated this observation using a patient-derived xenograft (PDX) obtained from a human breast tumor. Cells derived from this PDX were treated either in the absence or presence of IKE for 14 days and FRα expression was assessed by flow cytometry. Cells that survived IKE treatment exhibited significantly more FRα expression than control cells (**Fig. 1B**).

**Figure 1:**
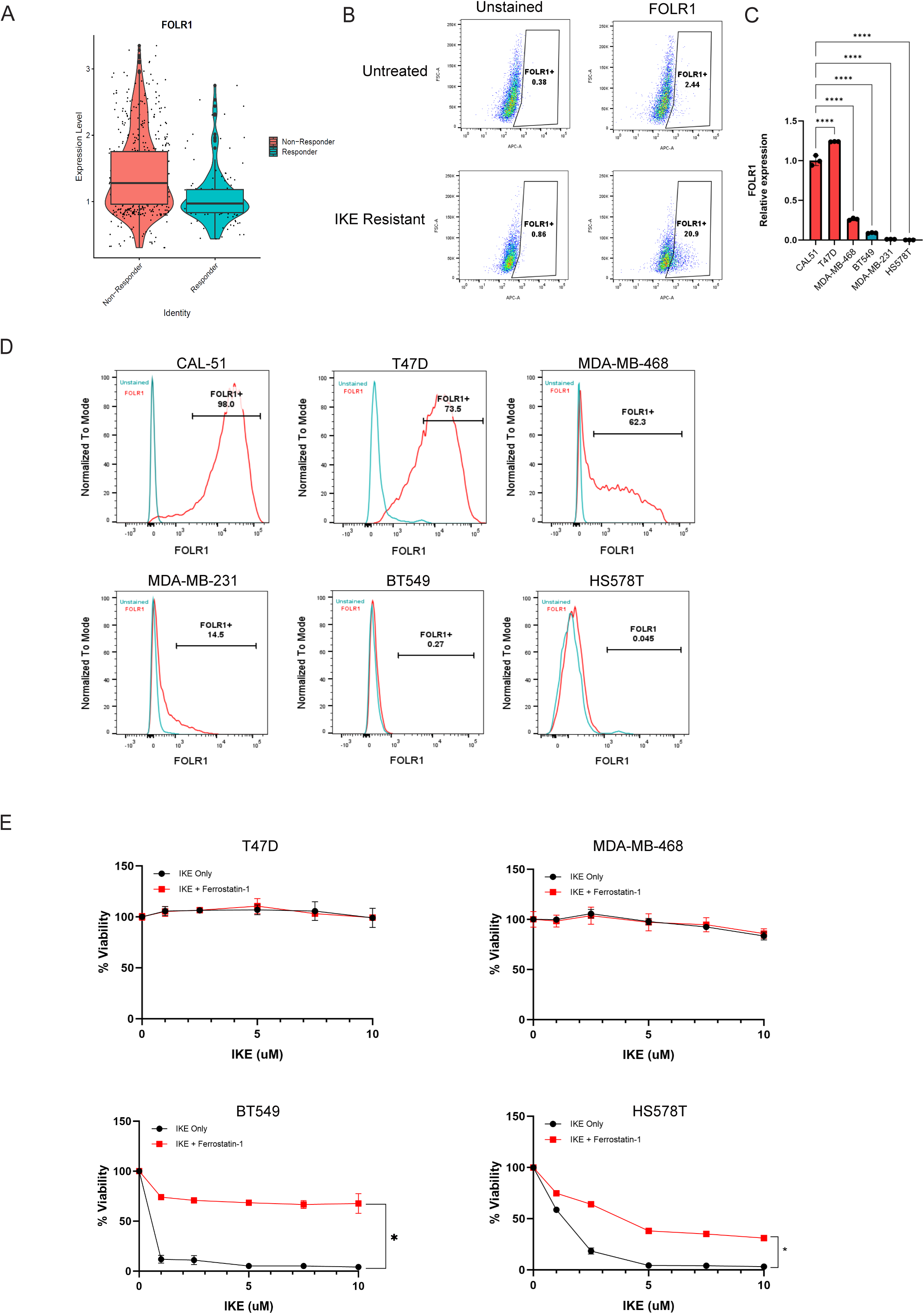
FRα expression correlates with ferroptosis resistance in breast cancer cell lines. (A) Expression of FOLR1 in IKE responder and non-responder cell populations obtained from scRNA sequencing analysis. (B) Primary tumors from PDX model were isolated and either untreated or treated with 1 uM IKE for 2 weeks. Flow cytometry of 5000 events for surface expression of folate receptor α (FRα) were assessed (C) Bar graphs show fold change of FOLR1/GAPDH mRNA in a panel of six breast cancer cell lines measured by qPCR (n = 3 replicates). Statistical significance was determined by one-way ANOVA, ****p < 0.0001 (D) Flow cytometry captured 10,000 events of cells for folate receptor α (FRα) surface expression in 6 breast cancer cell lines (n = 2 replicates) (E) T47D, MDA-MB-468, BT549 and Hs578T cell lines treated with increasing concentrations of IKE alone or in combination with ferrostatin-1 (2 uM) (n = 3 replicates). Cells were assessed for viability by Cell Titer Glo after 48 h. Error bars represent SD. Statistical significance was determined by 2-sided, unpaired t test, *p<0.05.

Subsequently, we investigated whether the expression of folate receptor-α (FRα) correlates with resistance to IKE in breast cancer cell lines. We analyzed 6 cell lines (CAL-51, T47D, MDA-MB-468, BT549, MDA-MB-231, Hs578T) that had been characterized previously for FRα mRNA expression [9] and verified that its expression is significantly higher in CAL-51, T47D and MDA-MB-468 cells compared to BT549, MDA-MB-231 and Hs578T cells (**Fig. 1C**). We also extended this analysis to validate surface expression of FRα as assessed by flow cytometry (**Fig. 1D**). Importantly, we observed a direct correlation between FRα surface expression and resistance to IKE. CAL-51[14], MDA-MB-468 and T47D (**Fig. 1D**) cells were resistant to IKE treatment (**Fig. 1E**). In contrast, MDA-MB-231[14], BT549 and Hs578T cells were sensitive to IKE, with the ferroptosis inhibitor, ferrostatin-1, rescuing loss of viability (**Fig. 1E**).

### Folate receptor α has a causal role in promoting ferroptosis resistance

To determine whether FRα has a causal role in ferroptosis sensitivity, we used two independent shRNAs to diminish its expression in CAL-51 (**Fig. 2A**) and T47D cells (**Fig. 2D**), cell lines that are resistant to IKE (**Fig. 1E**) and have high FRα surface expression (**Fig. 1D**). Diminished FRα expression decreased the ability of both cell lines to survive ECM-detachment compared to parental control cells, an effect that was rescued by ferrostatin-1 (**Fig. 2B, E**). In addition, FRα knockdown cells were more susceptible to IKE than control cells, an effect that was also rescued by ferrostatin-1 (**Fig. 2C, 2F)**.

**Figure 2:**
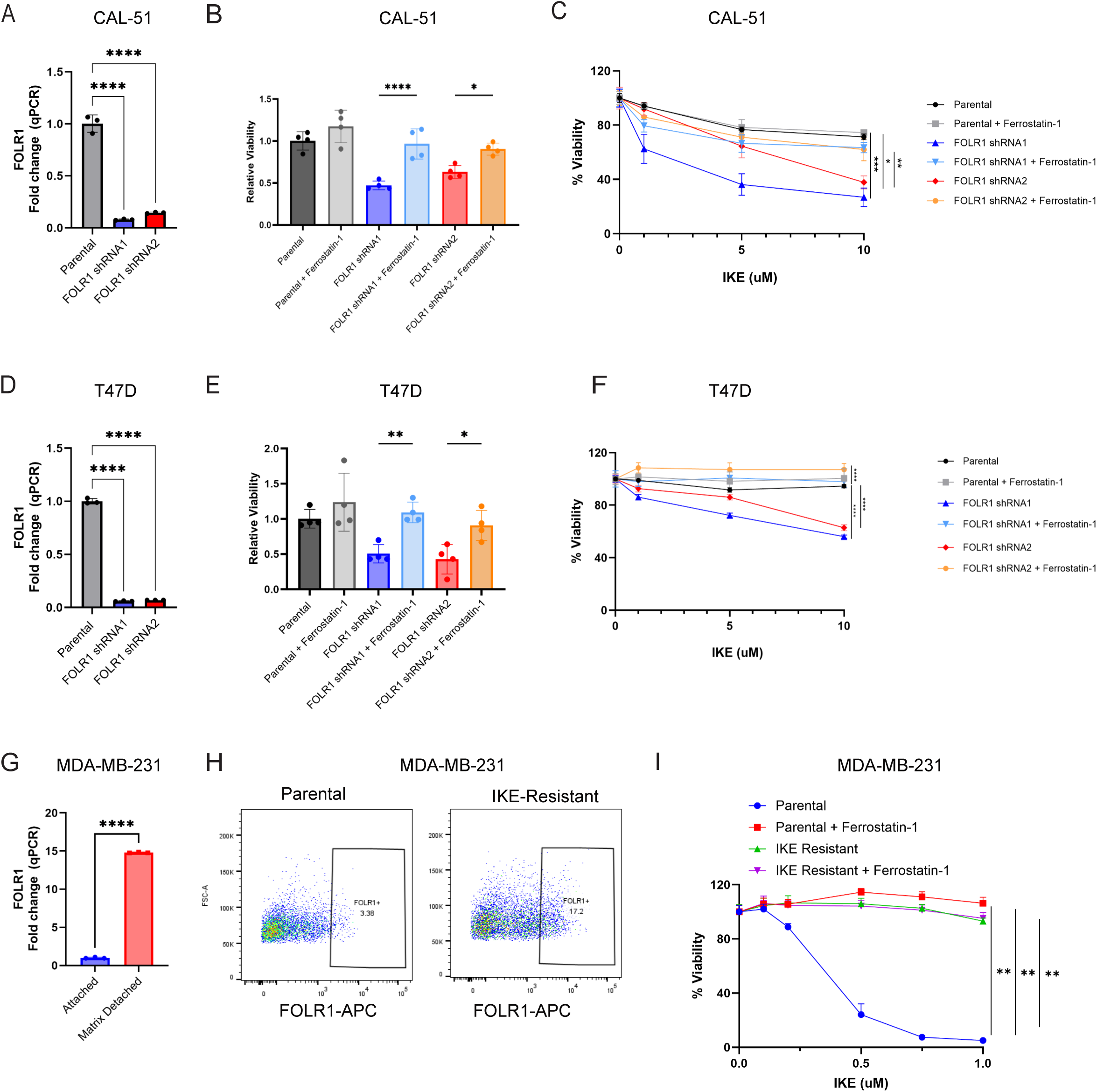
FRα has a causal role in ferroptosis resistance. (A) Validation of FOLR1 knockdown in CAL-51 cells transduced with FOLR1 shRNAs (shRNA1 and shRNA2) compared with parental cells by qPCR (n = 3 replicates) Statistical significance was determined by one-way ANOVA, ***p < 0.0001 (B) Matrix detachment of parental CAL-51 and FOLR1 knockdown cells performed in presence of either DMSO or ferrostatin-1 (2 uM) and cell viability was quantified over 48 hr (n = 3 biological replicates). Statistical significance was determined by one-way ANOVA, *p<0.05, **p < 0.01, ***p<0.001. Error bars represent SD. (C) Parental CAL-51 and respective FOLR1 shRNA1 and FOLR1 shRNA2 cells were treated with range of IKE concentrations alone or in combination with ferrostatin-1 (2uM). Cell viability was assessed after 48h. Statistical significance was determined by 2-way ANOVA, *p<0.05, ****p < 0.0001. Data are means ± SD of 3 biological replicates (D) Validation of FOLR1 knockdown in T47D cells transduced with FOLR1 shRNAs (shRNA1 and shRNA2) compared with parental cells by qPCR (n = 3 replicates). Statistical significance was determined by one-way ANOVA, ***p < 0.0001 (E) Matrix detachment of parental T47D and FOLR1 knockdown cells in presence of DMSO or ferrostatin-1 (2 uM) and cell viability was quantified over 48 hr (n = 4 biological replicates). Statistical significance was determined by one-way ANOVA, *p<0.05, **p < 0.01, ***p<0.001. Error bars represent SD. (F) Parental T47D, FOLR1 shRNA1 and FOLR1 shRNA2 cells were treated with range of IKE concentrations alone or in combination with ferrostatin-1 (2uM). Cell viability was assessed after 48h. Statistical significance was determined by 2-way ANOVA, *p<0.05, ****p < 0.0001. Data are means ± SD of two biological replicates. (G) MDA-MB-231 cells were cultured in attached or matrix detached conditions for 72 hrs. FOLR1 mRNA expression was compared between conditions by qPCR. (H) FOLR1 surface expression was measured between parental and IKE-Resistant MDA-MB-231 cells by flow cytometry. 5000 events were recorded. (n=2 biological replicates). (I) Parental and IKE-Resistant MDA-MB-231 cells were treated with range of IKE concentrations alone or in combination with ferrostatin-1 (2uM). Cell viability was assessed after 48h. Statistical significance was determined by 2-way ANOVA, *p<0.05, ****p < 0.0001. Data are means ± SD of three biological replicates.

We also investigated whether cells selected for ferroptosis resistance exhibited enhanced FRα expression. Although adherent MDA-MB-231 cells are sensitive to ferroptosis-inducing drugs and express low levels of FRα (**Fig. 1C**), we observed that ECM detachment markedly increased FRα expression in these cells (**Fig. 2G**), consistent with the reported ferroptosis resistance of these cells under detached conditions[15, 16]. We also observed that long-term IKE treatment of these cells induced FRα expression in a subpopulation of cells **(Fig. 2H)**. This FRα-expressing subpopulation was sufficient to protect the bulk IKE-resistant population from ferroptosis, as assessed by cell viability following IKE treatment **(Fig. 2I)**.

### FRα signaling promotes ferroptosis resistance by activating STAT3

To identify the signaling pathway that promotes ferroptosis resistance downstream of FRα, we performed GSEA on our scRNA-seq data to compare the transcriptomes of IKE responder and non-responder populations. This analysis revealed enrichment of a JAK-STAT gene set in the IKE non-responder compared to responder populations **(Fig. 3A)**. This result captured our attention for two reasons. First, ECM detachment can activate STAT3[17]. Second, non-canonical FRα signaling is known to activate JAK/STAT3 [15–19], which can also contribute to ferroptosis resistance [20, 21]. For these reasons, we focused on FRα-mediated JAK/STAT3 activation. To confirm the involvement of JAK kinase in the activation of STAT3, we treated parental CAL-51 cells with the JAK inhibitor, ruxolitinib, which decreased STAT3 phosphorylation **(Fig. 3B)**. In support of the hypothesis that FRα mediates STAT3 activation, we verified that FRα contributes to STAT3 phosphorylation in both CAL-51 (**Fig. 3D**) and T47D (**Fig. 3E**) cells. We also observed increased STAT3 phosphorylation in IKE resistant MDA-MB-231 compared to the parental cell line **(Fig. 3F)** and that knocking down STAT3 expression (**Fig. 3G, H**) sensitized both cell lines to IKE (**Fig. 3I, J**). Importantly, we also demonstrated that the ability of FRα to promote ferroptosis resistance is dependent on STAT3. Specifically, we observed that expression of constitutively active (CA) STAT3 **(Fig. S2A**) was able to rescue resistance to IKE in FRα-knockdown cells (**Fig. 3K**).

**Figure 3:**
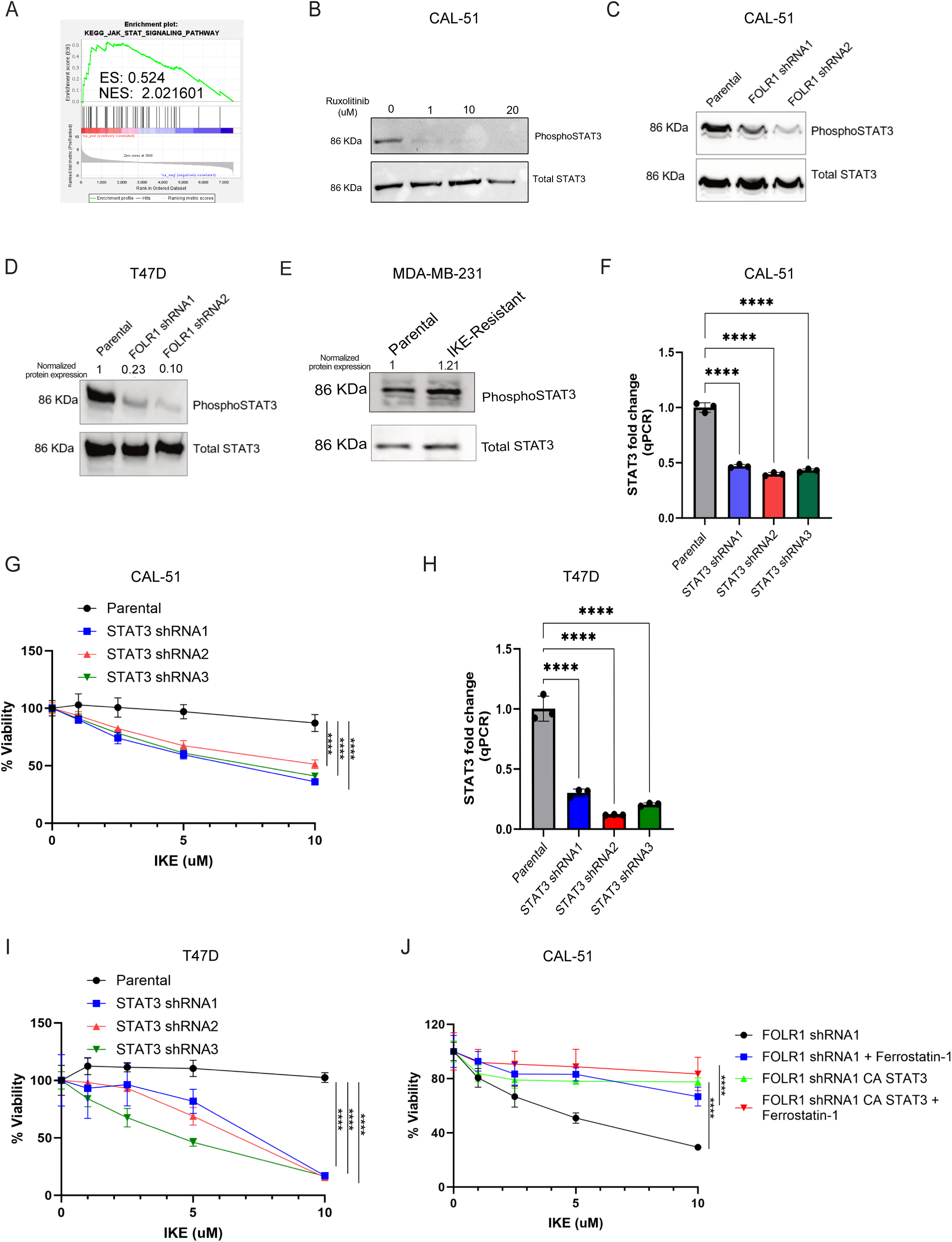
FRα activation of STAT3 mediates ferroptosis resistance. (A) GSEA analysis shows enrichment of JAK-STAT pathway in IKE non-responder population compared to responders from scRNA-seq dataset. (B) Phosphorylated STAT3-Tyr705 in parental CAL-51 upon treatment with DMSO or Ruxolitinib (1 uM, 10 uM, 20 uM) measured by immunoblotting (n=1). Total STAT3 was used as a loading control here. (C) Expression of STAT3 in IKE Responder and Non-Responder cell populations obtained from scRNA sequencing analysis. (D) Phosphorylated STAT3-Tyr705 was compared in parental CAL-51 with respective FOLR1 knockdown cells by immunoblotting (n = 3 biological replicates). Total STAT3 was used as a loading control. (E) Phosphorylated STAT3-Tyr705 was compared in parental T47D cells with respective FOLR1 knockdown cells by immunoblotting (n = 2 replicates). Total STAT3 was used as a loading control. (F) Phosphorylated STAT3-Tyr705 was compared in parental and IKE-Resistant MDA-MB-231 cells by western blotting (n=2 replicates). Total STAT3 was used as a loading control. (G, H) STAT3 knockdown was measured in CAL-51 and T47D cells transduced with STAT3 shRNAs (shRNA1, shRNA2, shRNA3) relative to parental cells by qPCR (n = 3 replicates). Statistical significance was determined by one-way ANOVA, ****p <0.0001. (I, J) Parental CAL-51 or T47D cells, respective STAT3 knockdown cell lines i.e. STAT3 shRNA1, STAT3 shRNA2 and STAT3 shRNA3 cells treated with range of IKE concentration (n = 2 replicates). Cell viability was assessed after 48h. (K) CAL-51 FOLR1 shRNA1 cells were transduced with a construct expressing constitutively active STAT3 and treated with increasing concentrations of IKE alone or in combination with ferrostatin-1 rescue (2 uM) (n = 3 replicates). Cell viability was assessed after 48h. Error bars represent SD. Statistical significance was determined by 2-way ANOVA. *p<0.05, **p < 0.01, ***p<0.001, ****p < 0.0001.

### FRα-mediated activation of STAT3 induces the transcription of LATS1 to inhibit YAP

In pursuit of the mechanism by which FRα-mediated STAT3 activation promotes ferroptosis resistance, we were drawn to a report that ECM detachment of carcinoma cells activates Hippo signaling by inducing LATS1 [22]. This finding captured our attention because there is considerable evidence, including from our own lab, that Hippo signaling promotes resistance to ferroptosis [8, 23, 24]. To assess this possibility, we compared *LATS1* mRNA expression in adherent versus ECM-detached CAL-51 cells and observed a significant increase in *LATS1* expression in ECM-detached cells **(Fig. 4A).** Subsequently, we evaluated the ability of FRα to regulate *LATS1* mRNA expression. Knockdown of FRα in both CAL-51 (**Fig. 4B**) and T47D cells (**Fig. 4C**) resulted in a decrease in *LATS1* mRNA expression. Moreover, IKE resistant MDA-MB-231 cells also showed increased *LATS1* mRNA expression compared to parental control **(Fig. 4D)**. Next, we evaluated the ability of STAT3 to regulate *LATS1* expression. Knockdown of STAT3 in CAL-51 cells decreased *LATS1* mRNA expression (**Fig. 4E)** and the treatment of these cells with a specific STAT3 inhibitor (Stattic) decreased *LATS1* mRNA (**Fig. 4F**) and protein (**Fig. 4G**) expression. Similar results were obtained with T47D cells (**Fig. 4H, 4I**). Together, these findings indicate that FRα-mediated STAT3 activation regulates LATS1 expression. Because STAT3 often functions as a transcription factor, we hypothesized that it regulates *LATS1* transcription. This hypothesis was strengthened by our observation that the *LATS1* promoter contains a canonical STAT3 binding site (**Fig. 4J**) [25], and we confirmed by ChIP that phospho-STAT3 binds to this site (**Fig. 4K**). Also, treating CAL-51 cells with a LATS1/2 inhibitor sensitized parental cells to IKE, an effect which was rescued by ferrostatin-1, confirming that LATS1 signaling mediates ferroptosis resistance (**Fig. 4L**).

**Figure 4:**
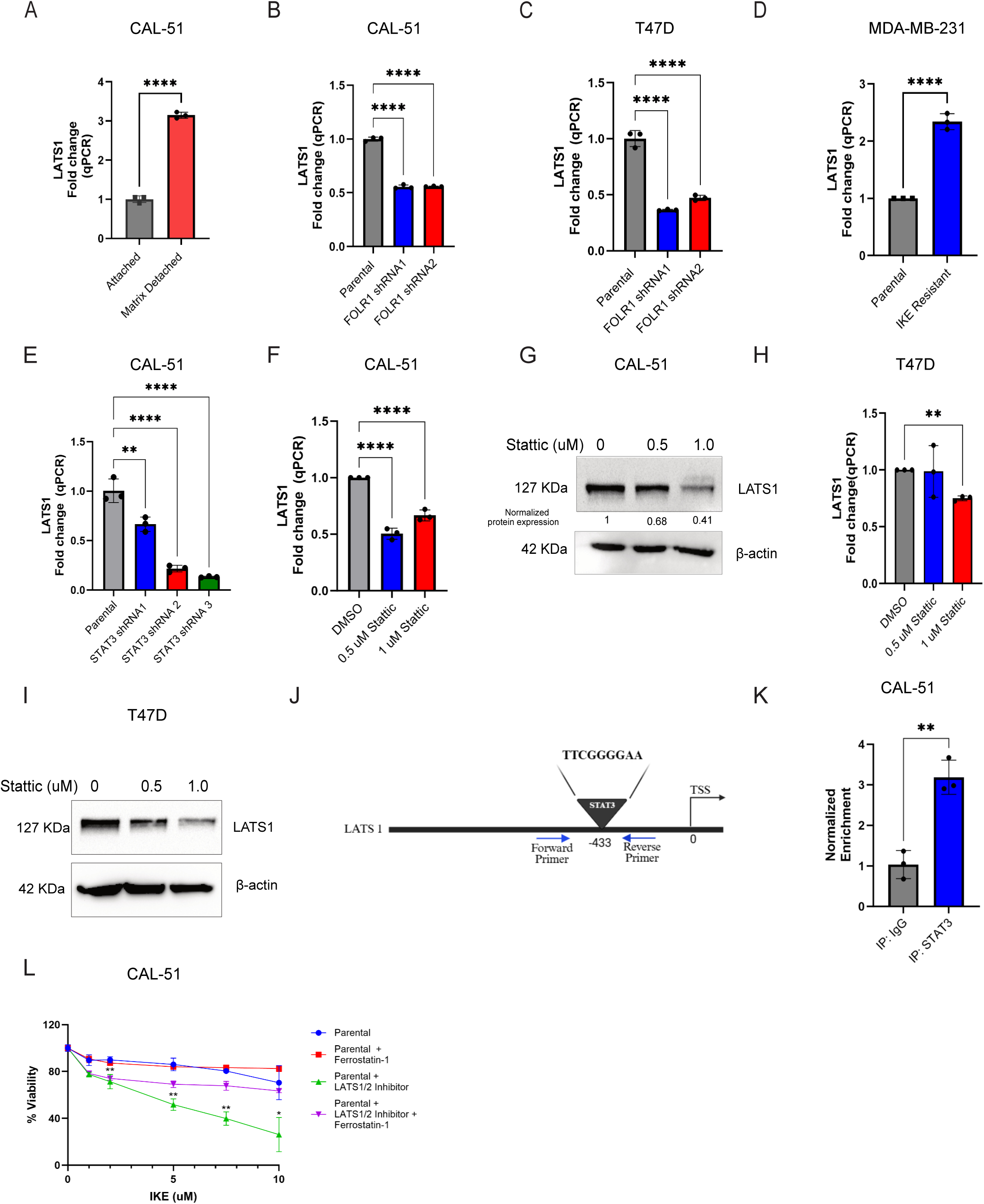
FRα-mediated STAT3 activation induces LATS1. (A) CAL-51 cells were maintained in either attached or matrix-detached conditions for 48 hrs. LATS1 mRNA expression was measured by qPCR. (B, C) LATS1 mRNA expression measured in parental CAL-51, T47D and FOLR1 knockdown cells by qPCR (n=3 replicates) (D) LATS1 mRNA expression measured between parental and IKE-Resistant MDA-MB-231 cells by qPCR. (E) LATS1 mRNA expression quantified in parental CAL-51 and STAT3 knockdown cells by qPCR (n=3 replicates) (F, G) LATS1 mRNA and protein expression compared between parental CAL-51 treated with DMSO and STAT3 inhibitor stattic (0.5 uM, 1 uM) for 48 h (n=2 replicates). β-actin was used as a loading control. (H, I) LATS1 mRNA and protein expression compared between parental T47D treated with DMSO and STAT3 inhibitor stattic (0.5 uM, 1 uM) for 48 h (n=2 replicates). β-actin was used as a loading control. (J) Schematic showing the position of primer pairs and putative STAT3 DNA motif on LATS1 promoter region. (K) Chromatin from CAL-51 cells was immunoprecipitated with STAT3 or IgG antibody and analyzed by qPCR using primers specific to putative STAT3 binding site on LATS1 promoter (n=3 replicates). Statistical significance was determined by unpaired t-test, **p < 0.01. (L) Parental CAL-51 cells were pretreated with 1 μM of LATS1/2 inhibitor (TRULI) for 1hour followed by treatment with either a DMSO vehicle control or range of IKE concentration (1 μM,2 μM,5 μM,7.5 μM,10 μM) with a 2μM ferrostatin-1 rescue for and cell viability was assessed by CTG assay after 48 hr. Statistical significance was determined by 2-way ANOVA, *p<0.05, **p<0.01. (n= 4 replicates) Error bars represent SD. Statistical significance was determined by one-way ANOVA. *p<0.05, **p<0.01, ***p<0.001, ****p < 0.0001.

Based on these observations, we hypothesized that FRα-mediated STAT3 activation promotes ferroptosis resistance, in part, by inducing LATS1 and thereby inhibiting YAP, which can sensitize cells to ferroptosis[8, 23, 24]. In support of this hypothesis, we observed that the expression of canonical YAP target genes (*CTGF* and *ANKRD1*) was significantly downregulated in ECM-detached CAL-51 cells compared to adherent cells **(Fig. 5A).** We also observed that knocking down FRα in CAL-51 (**Fig. 5B**) and T47D cells (**Fig. 5C**) increased the expression of *CTGF* and *ANKRD1* and increased YAP nuclear localization (**Fig. 5D**). To verify a causal role for YAP in ferroptosis sensitivity, we treated FRα knockdown CAL-51 cells with the YAP inhibitor verteporfin and observed that it decreased the sensitivity of these cells to IKE compared to control cells **(Fig. 5E)**. Consistent with our hypothesis, knockdown of STAT3 in CAL-51 and T47D cells also increased the expression of *CTGF*, CYR61 and *ANKRD1* (**Fig. 5F, 5G**). We also observed that expression of CA STAT3 in FRα-knockdown cells decreased the expression of YAP target genes *(CTGF and CYR61)* **(Fig. 5H, 5I)**, supporting the hypothesis that FRα-mediated repression of YAP is STAT3-dependent.

**Figure 5:**
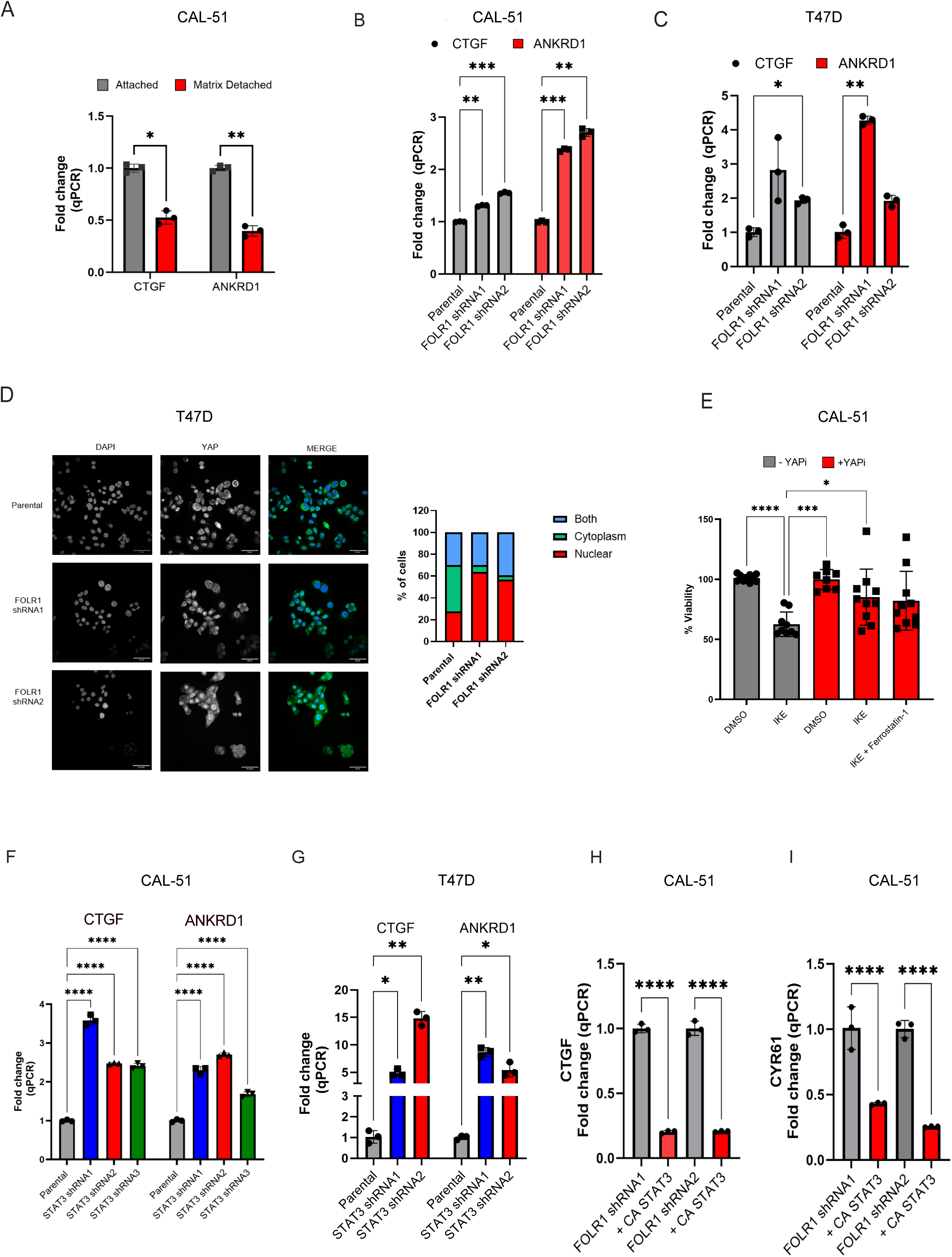
FRα-mediated STAT3 activation inhibits YAP activation. (A) CAL-51 cells were maintained in either attached or matrix-detached conditions for 48 hrs. YAP target genes (CTGF and ANKRD1) mRNA expression was measured by qPCR. (B) YAP target genes (CTGF and ANKRD1) mRNA expression in parental CAL-51 with FOLR1 shRNA cell lines quantified by qPCR (n =3 replicates) (C) YAP target genes (CTGF and ANKRD1) mRNA expression in parental T47D and FOLR1 shRNA cell lines quantified by qPCR (n =3 replicates) (D) Representative immunofluorescence images of YAP and DAPI staining in parental T47D and FOLR1 knockdown cells with quantification of YAP localization in nuclear, cytoplasmic or both regions (n=3 replicates). Scale bars: 50 um. (E) FRα knockdown CAL-51 cells were pretreated with 10 μM of YAP inhibitor (verteporfin) 24 hours followed by treatment with either a DMSO vehicle control, 5 μM IKE, or IKE with a 2μM ferrostatin-1 rescue for another 24 hours. Cell viability was assessed by CTG assay. Statistical significance was determined by unpaired t-test, ***p < 0.001. (n= 3 biological replicates) (F, G) CTGF and ANKRD1 mRNA expression in parental CAL-51 and T47D and STAT3 knockdown cells assessed by qPCR (n=3 replicates) (H, I) CTGF and CYR61 mRNA expression compared between CAL-51 FOLR1 knockdown and FOLR1 knockdown cells with restored constitutively active STAT3 (CA STAT3) measured by qPCR (n=3 replicates). Statistical significance was determined by one-way ANOVA, ****p < 0.0001. Error bars represent SD. Statistical significance was determined by 2-way ANOVA. *p<0.05, **p < 0.01, ***p<0.001, ****p < 0.0001.

### FRα promotes ferroptosis resistance by STAT3/LATS1-mediated YAP inhibition and suppression of ACSL4

To investigate how inhibition of YAP by the FRα/STAT3/LATS1 axis contributes to ferroptosis resistance, we focused on the ability of YAP to regulate acyl-CoA synthetase long-chain family member 4 (ACSL4), a metabolic enzyme that is an essential component of the ferroptotic process[26]. This enzyme catalyzes the transformation of PUFAs into phospholipid PUFAs (PL-PUFAs), which are incorporated into plasma membranes and act as the primary substrates for lipid peroxidation. Previous work has shown that ACSL4 is a YAP transcriptional target through direct binding of YAP to the ACSL4 promoter [8, 27], implicating YAP in the regulation of ACSL4 expression as a mechanism by which it promotes ferroptosis sensitivity. For these reasons, we investigated whether the FRα/STAT3/LATS1 axis suppresses ACSL4. Using the CCLE database, we found an inverse correlation between FOLR1 and ACSL4 expression in breast cancer cell lines **(Fig. 6A)**. Knockdown of FRα increased ACSL4 protein and mRNA expression in CAL-51 (**Fig. 6B, 6C)** and T47D (**Fig. 6D, E**) cells, and we observed decreased expression of ACSL4 in IKE resistant MDA-MB-231 cells compared to parental cells **(Fig. 6F)**. Also, inhibition of STAT3 expression using shRNAs increased ACSL4 mRNA and protein expression in CAL-51 and T47D cells (**Fig. 6G, 6I and 6J**). Moreover, expression of CA STAT3 in FRα-knockdown cells decreased the expression of ACSL4 **(Fig. 6H)**, supporting the hypothesis that FRα-mediated repression of ACSL4 is STAT3-dependent. Furthermore, a LATS1/2 inhibitor increased ACSL4 expression in a dose-dependent manner in parental CAL-51 cells **(Fig. 6K)**. Lastly, CAL-51 FRα knockdown cells exhibited increased lipid peroxidation, a potential consequence of increased ACSL4 activity, compared to control cells upon IKE treatment **(Fig. 6L)**.

**Figure 6:**
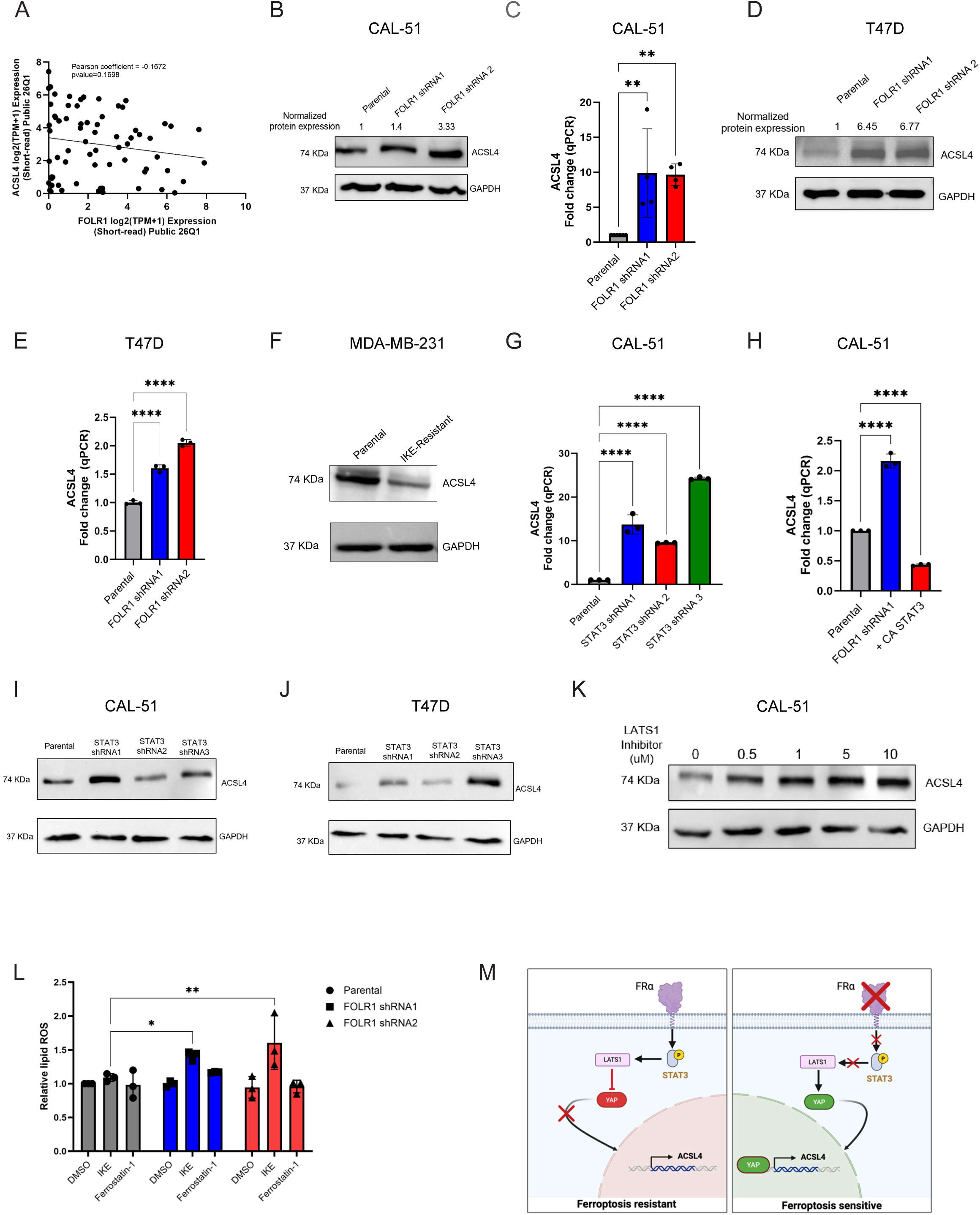
FRα-mediated STAT3 activation represses ACSL4 to mediate ferroptosis resistance. (A) Graph depicting a correlation curve between ACSL4 and FOLR1 gene expression obtained from DepMap database for breast cancer cell lines. Correlation coefficients and statistical significance were obtained by analyzing Pearson’s correlation on GraphPad Prism. (B) ACSL4 protein level was assessed by immunoblotting in parental CAL-51 and FOLR1 knockdown cells (FOLR1 shRNA1 and FOLR1 shRNA2) (n=2 replicates). GAPDH was used as a loading control. (C) ACSL4 mRNA expression was assessed by qPCR in parental CAL-51 and FOLR1 knockdown cells (FOLR1 shRNA1 and FOLR1 shRNA2) (n=3 replicates). (D) ACSL4 protein level compared between parental T47D and FOLR1 knockdown cells (FOLR1 shRNA1 and FOLR1 shRNA2) by immunoblotting (n=3 replicates) (E) ACSL4 mRNA expression was assessed by qPCR in parental T47D and FOLR1 knockdown cells (FOLR1 shRNA1 and FOLR1 shRNA2) (n=3 replicates). (F) ACSL4 protein expression was measured between parental and IKE-Resistant MDA-MB-231 cells by immunoblotting. GAPDH was used as a loading control. (G) Bar graphs showing fold change of ACSL4 mRNA in parental CAL-51 and STAT3 knockdown cells (STAT3 shRNA1, STAT3 shRNA2, STAT3 shRNA3) by qPCR (n=3 replicates). (H) ACSL4 mRNA expression measured by qPCR in parental CAL-51, CAL-51 FOLR1 shRNA1 and FOLR1 shRNA1 CA STAT3 cells (n=3 replicates). (I) ACSL4 protein expression measured between parental CAL-51 and STAT3 knockdown cells (STAT3 shRNA1, STAT3 shRNA2, STAT3 shRNA3) by immunoblotting (n=2 biological replicates). GAPDH was used as a loading control. (J) ACSL4 protein expression measured between parental T47D and STAT3 knockdown cells (STAT3 shRNA1, STAT3 shRNA2, STAT3 shRNA3) by immunoblotting (n=2 biological replicates). GAPDH was used as a loading control (K) CAL-51 cells were treated with LATS1/2 inhibitor (TRULI) at 0, 0.5, 1, 5 and 10 uM for 48 hrs. ACSL4 protein expression was measured by immunoblotting. GAPDH was used as a loading control. (L) C11 bodipy staining of parental CAL-51, FOLR1 shRNA1 and FOLR1 shRNA2 cells treated with DMSO or IKE (10 uM) for 24 hr (n=3 biological replicates). (M) Schematic of the proposed mechanism by which FRα mediates ferroptosis resistance. Breast cancer cells expressing FRα mediate ferroptosis resistance by its non-canonical activation of STAT3 that regulates the transcription of LATS1 and the consequent inhibition of the Hippo effector, YAP. YAP inhibition decreases the expression of ACSL4, a critical effector of ferroptosis that generates the PL-PUFAs that are the target of lipid peroxidation. Error bars represent SD. Statistical significance was determined by unpaired t-test or 2-way ANOVA. *p<0.05, **p < 0.01, ***p<0.001, ****p < 0.0001.

## Discussion

The data reported in this study reveal a novel role for FRα in enabling cancer cells to survive the ferroptotic stress of ECM detachment, a process that is relevant to the conditions encountered by metastatic cells [17], as well as in response to the ferroptosis-inducing drug IKE. Our data also highlight the involvement of FRα-mediated activation of STAT3 as a core mechanism of ferroptosis resistance. We extend this observation by identifying a novel STAT3-mediated ferroptosis resistance mechanism that regulates transcription of the Hippo inhibitor *LATS1*, whose expression is also induced upon ECM detachment. Consequently, LATS1 represses YAP-mediated regulation of ACSL4, a critical effector of ferroptosis that generates the PL-PUFAs that are the target of lipid peroxidation. Thus, our findings, which are schematized in **Fig. 6M**, implicate FRα in the mitigation of lipid peroxidation, which is the root cause of ferroptosis.

A strength of our study is that we associated FRα with ferroptosis resistance based on an unbiased bioinformatics analysis of scRNA-seq data on ferroptosis sensitive and resistant populations in breast cancer organoids. Although several surface proteins including the reduced folate carrier and the proton-coupled folate transporter can transport folate into cells in addition to FRα [16], they were not identified in our analysis of ferroptosis-resistant populations. This observation is consistent with our conclusion that the ability of FRα to promote ferroptosis resistance is mediated by a non-canonical signaling mechanism that involves the ability of FRα to activate STAT3. Indeed, there is compelling evidence in the literature for such non-canonical signaling, especially in tumor cells [18, 19], which is substantiated by the data we provide.

Our data implicate LATS1/2 as critical effectors of ferroptosis resistance because of their ability to inhibit YAP and YAP-mediated regulation of ACSL4. This finding builds on the report that ECM-detachment induces LATS1/2 resulting in YAP phosphorylation and inhibition[20]. This study, however, did not implicate LATS1/2-mediated regulation of YAP in ferroptosis resistance. We also demonstrated that FRα-mediated STAT3 activation has a key role in this process because STAT3 regulates *LATS1* transcription. This result provides one mechanism by which the activity of YAP can be regulated to affect ferroptosis. It also reveals a novel transcriptional target of STAT3 that contributes to ferroptosis resistance. We note, however, that other STAT3 transcriptional targets including *SLC7A11* and *GPX4* contribute to ferroptosis resistance [21], and we emphasize that our data do not exclude the potential contribution of these other targets to ferroptosis resistance in our models.

We focused on the ability of breast cancer cells to survive the ferroptotic stress imposed by ECM-detachment for several reasons. This survival mechanism, often referred to as anoikis, has long been thought to depend solely on the ability of cells to resist apoptosis [22]. More recent studies by us and others, however, have revealed that the ability to evade ferroptosis is necessary for anoikis because this process generates a pro-ferroptotic environment by increasing the concentration of intracellular ROS [4, 23, 24]. Moreover, our finding that FRα enables breast cancer cells to survive ECM-detachment by resisting ferroptosis is highly significant because the ability of cells to survive under these conditions can be prognostic for their metastatic potential[6]. Although ferroptosis resistance is most often defined as the ability to resist drugs that trigger ferroptosis such as IKE, which we demonstrated in this study, we believe that the ability of tumor cells to resist physiological conditions that they encounter during their progression is a more relevant definition in the context of mechanistic cancer biology. This consideration is reinforced by the reports that the ability to evade ferroptosis is an essential component of metastasis[25–27].

Our results also bear on other studies that investigated the impact of ECM detachment and 3D culture conditions on ferroptosis resistance. Recently, it was reported that ECM detached tumor cells are more ferroptosis resistant than the same cells maintained under adherent conditions[15]. This study concluded that the expression of the monounsaturated fatty acid (MUFA) biosynthetic gene stearoyl CoA desaturase, which alters the ratio of MUFA-phospholipids to PUFA-phospholipids to favor ferroptosis resistance, increased in response to ECM detachment. We also observed that ECM-detached CAL-51 and T47D cells can survive ECM detachment and that FRα has a causal role in this process. Our results are compatible with the previous study, because we conclude that FRα mediates ferroptosis resistance by repressing the expression of ACSL4, which promotes the incorporation of oxidizable PUFAs into membrane phospholipids, and that this effect is amplified in 3D culture. Although FRα does not appear to regulate stearoyl CoA desaturase in our models (**Supp Fig. 4**), it does impact lipid metabolism by regulating ACSL4 to affect ferroptosis resistance. Moreover, we provide a mechanism for how a cell surface receptor, whose expression is regulated by ECM detachment, can regulate the expression of a key enzyme involved in lipid metabolism.

Our results also intersect closely with a study that investigated the mechanistic basis of ferroptosis resistance in ECM-detached carcinoma cells, including MDA-MB-231 cells [16]. Like us, that study found that ECM detachment reduces YAP-mediated ACSL4 expression and lowers PUFA-containing membrane phospholipids. However, restoring ACSL4 with a phosphorylation-resistant YAP mutant (YAP S127A) was not enough by itself to rescue resistance to erastin- or RSL3-induced ferroptosis. Instead, the authors identified an NRF2-dependent iron-reprogramming pathway involving reduced transferrin receptor 1-mediated iron uptake and increased ferritin (FTH1/FTL) storage as the main driver of resistance in that setting. We think the two mechanisms are complementary rather than contradictory because they could act in parallel: ferroptotic lipid peroxidation requires both oxidizable PUFA-PL substrates and redox-active iron, so limiting either one can confer resistance. Whether FRα-mediated STAT3 signaling also affects iron handling independently of ACSL4 needs further investigation. Together, the two studies show that ECM-detached cells can draw on multiple, non-exclusive resistance mechanisms, underscoring the complexity of this survival strategy during metastasis.

It is important to discuss our findings in the context of other studies that investigated FRα in breast cancer, especially TNBC[11]. The conclusion from this work is that FRα is expressed preferentially in TNBC compared to other breast cancer subtypes and that it contributes to tumor growth. We do not challenge these findings specifically, but our data reveal the heterogeneity of FRα expression even within TNBC and that its expression is influenced significantly by physiological conditions. A distinction of our study is that we used scRNA-seq to discover that FRα expression is elevated in distinct sub-populations of TNBC organoids that were identified by their resistance to IKE. We suggest that ferroptotic stress either selects for the survival of FRα-expressing sub-populations or induces FRα expression. This conclusion is justified by our findings that cells derived from a TNBC PDX that survive IKE treatment exhibit much higher FRα expression than control cells (**Fig. 1B**) and that ECM detachment of MDA-MB-231 cells increases FRα expression markedly (**Fig. 2G**).

In summary, we describe a novel function for FRα in the ability of breast cancer cells to resist the ferroptotic stress imposed by ECM detachment, as well as IKE treatment, that is dependent on its non-canonical signaling function that is intimately associated with Hippo signaling. Implicit in our data is the plasticity of FRα expression in breast cancer that is linked to a more aggressive and potentially metastatic phenotype. These data provide a rationale for future studies on its contribution to metastasis and therapeutic potential.

## Materials and Methods

### Cell lines

The following cell lines were used in this study: CAL-51 (human, female), T47D (human, female), MDA-MB-468 (human, female), MDA-MB-231 (human, female), BT549 (human, female), Hs578T (human, female). All cell lines were maintained at 37 ^°^C with 5% CO2 atmosphere in complete growth medium. Compounds used for treatment were added to and maintained in complete growth medium.

### Expression constructs

Lipofectamine 3000 (Thermo Fisher Scientific, cat. L3000008) was used for plasmid transfections according to manufacturer’s instructions. The folate receptor α plasmid was purchased from Origene (plasmid # OHu14848). After transfection, FRα^+^ cells were selected by Neomycin selection for 5-7 days and sorted via FACS. Sorted FRα^+^ cells were then used for downstream analysis such as gene expression and cell viability. The constitutively active STAT3 plasmid was purchased from Addgene (EF.STAT3C.Ubc.GFP plasmid #24983). Stable expression of constitutively active STAT3 in cells was obtained using a lentiviral vector. Following viral transduction with active STAT3 plasmid, cells were sorted for plasmid expression by a GFP selection marker.

### RNA Interference

Short hairpin RNAs were used to transiently knockdown expressions of FOLR1 and STAT3 in our cell lines using Lipofectamine 3000 (cat. L3000008). Stable expression of shRNA in cells was obtained using a lentiviral vector. The following shRNA constructs were obtained from UMass Chan RNAi Core: human FOLR1 (TRCN0000060343, TRCN0000060346) and human STAT3 (TRCN0000020840, TRCN0000020842, TRCN0000020843).

### Cell viability assay

Cells were seeded in a 96-well plate at 10,000 cells/well and treated with the DMSO, IKE or combination of IKE and ferrostatin-1 at described concentrations for 48 hrs. Cell viability was measured using Cell Titer-Glo (Promega, Cat no. G9242), where cells were incubated with Cell Titer-Glo reagent for 10 mins at room temperature according to the manufacturer’s instructions. Cell viability was assessed by measuring luminescence using a GloMax plate reader (Promega).

### Drug Treatment

Cells were seeded in 6-well plate and treated with inhibitors (Ruxolitinib/Stattic/LATS1/2 inhibitor-TRULI) for 48 hrs for immunoblotting with the exception of LATS1/2 inhibitor where cells were treated for 1 hour for PhosphoYAP expression.

### Matrix detachment assay

Matrix detachment assay was performed as described previously [6]. Briefly, 12- or 24-well plates were coated with poly 2-hydroxyethyl methacrylate (PolyHEMA, Sigma-Aldrich, 30mg/ml) in 95% EtOH and allowed to dry overnight. Cells were trypsinized, washed and resuspended in 0.2% Methylcellulose (MC) containing media to prevent cell clustering and subsequently placed onto Poly-HEMA coated plates. Treatments such as DMSO or Ferrostatin-1 were added to the MC media at the time of matrix detachment. Cell viability of matrix-detached cells was assessed by a hemocytometer using Trypan Blue exclusion for over 48 hours.

### Flow cytometry

To quantify surface expression of FRα, 200,000 cells were incubated in 100μL PBS containing FRα-APC conjugated antibody (Miltenyi Biotec, Cat no. 130-129-432) at 1:100 dilution. Cells were stained for 30 min on ice and then analyzed using a BioRad ZE5-A Cell Analyzer (BioRad). Median Fluorescence Intensity was calculated using FlowJo™ v10.8 software (BD Life Sciences).

### Quantitative real-time PCR

Total RNA extraction from cells was performed using Monarch® Spin RNA Isolation Kit [28] (Cat no. T2110S). cDNA synthesis was completed using Azura Genomics cDNA synthesis kit (Cat. AZ-1996). The relative expression levels were quantified using SYBR Green Master mix (Thermo Fisher, Cat. A46012). Experiments were performed in triplicate and normalized to GAPDH.

### Primers sequences

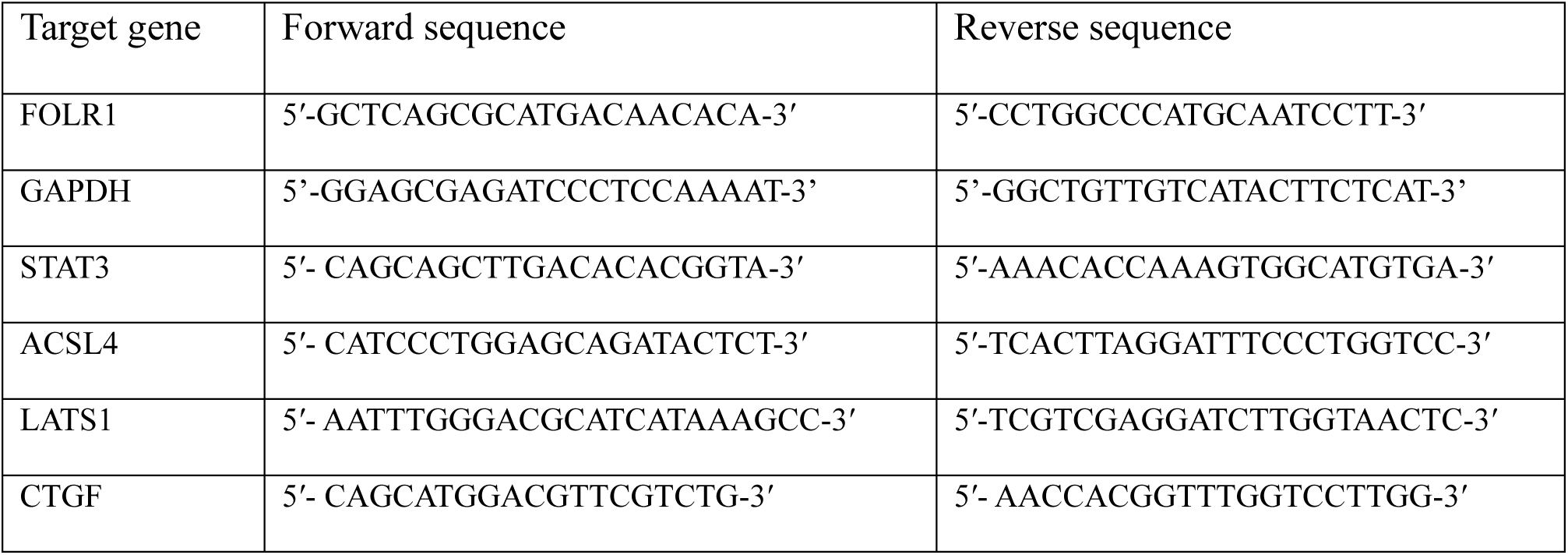
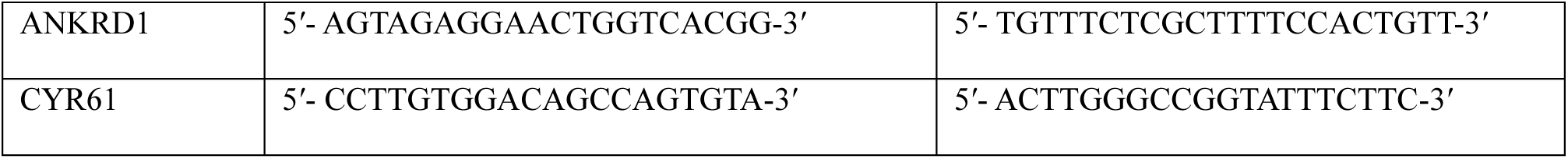

### scRNA seq

Processed single cell RNA seq data was downloaded from the Gene Expression Omnibus with the accession code GSE301773. Quality control, filtering, data normalization, variable extraction, data scaling, dimensionality reduction, and genetic imputation were performed as previously described [8]. Single Sample Gene Set Enrichment Analysis (ssGSEA) was performed using the escape package[29]. Unbiased clustering was performed using KNN (known nearest neighbors) clustering and differential expression was performed using the Seurat “Find Markers” function [30]. Cell clusters sensitive and resistant to IKE treatment were determined by comparing the median normalized ssGSEA enrichment score of a “response to erastin” gene signature which was determined by taking the top 50 differentially expressed genes from a previously published RNA seq dataset measuring the transcriptional response to erastin[31].

### ChIP

We used the ChIP-IT Express Chromatin Immunoprecipitation Kit (Active Motif) (Cat no. 53008). The following primers amplified the LATS1 promoter region with a STAT3 peak: LATS1 promoter region primer set: (forward, 5’-TCCTTCTACGGATGCGGTTG-3’; reverse, 5’-ACCTCCCTGTGGACCTCAGA-3’) Exon primer: (forward, 5’-CCTACACACCCTTCTTGGATAC-3’; reverse, 5’-TGGTGGTCCTT GATAGTTTGG-3’).

### Immunoblotting

Cells (2.5 x10^5^) were plated in a 6-well plate for 24 hrs. For protein extraction, cells were washed with cold PBS (1X). Cells were scraped on with 100 ul RIPA buffer (Sigma Aldrich, Cat no. R0278-50ML) containing 1X protease and phosphatase inhibitors (Thermo Fisher, Cat no. 78442). Protein concentration was determined by a colorimetric assay (Bio-Rad Protein Assay Dye Reagent Concentration, Cat no. 5000006). Subsequently, protein lysate was prepared in laemmli buffer (Boston Bioproducts, Cat no. BP-111R) by boiling for 5 mins at 95°C and separated on a 9% SDS-PAGE gel. Immunoblotting primary antibodies were used at the following concentrations:

P-YAP S127 (Cell signaling technology, Cat no. 4911S), 1:1,000; YAP D8H1X antibody (Cell signaling technology, cat no. 14074), 1:1,000, LATS1 (C66B5) (Cell signaling technology, Cat no. 3477), 1:1,000; GAPDH, (Cell signaling technology, Cat no. 2118S), 1:1,000; Phospho-Stat3 (Tyr705) (Cell signaling technology, Cat no. 9145), 1:1,000; streptavidin HRP (Cell signaling technology, Cat no. 7074S) 1:1,000.

### Immunofluorescence

Approximately 50,000 cells were plated on 35 mm ibidi slides (Ibidi, cat no. 81218–200). Supernatant was aspirated and cells were fixed with paraformaldehyde solution (4%) (Boston bioproducts, cat no. BM-155). Subsequently, cells were blocked using blocking buffer containing 1X PBS, 5% normal goat serum and 0.3% Triton X-100. Cells were incubated overnight at 4°C in YAP D8H1X antibody (Cell signaling technology, cat no. 14074) at 1:300 concentration in antibody dilution solution containing 1X PBS, 1% BSA and 0.3% Triton X-100. Next day, slides were washed with 1X PBS before incubation with either secondary antibody, or Hoechst stain (Invitrogen cat. A34055) for 1 hr at room temperature. Slides were mounted in 0.1 M n-propyl gallate, 90% (by volume) glycerol, and 10% PBS solution, and images were taken using a Nikon confocal microscope using a 60× oil objective. Images were processed using Fiji.

## Acknowledgments

We acknowledge Shivam Goel, a summer intern in the Mercurio lab who conceptualized this project and conducted the initial experiments that identified the folate receptor as a marker of ferroptosis resistance and established its functional role. We also thank Drs. Larry Matherly and Zhanjun Hou at the Karmanos Cancer Center for their valuable comments and advice.

---

**Supplementary Figure 1.**
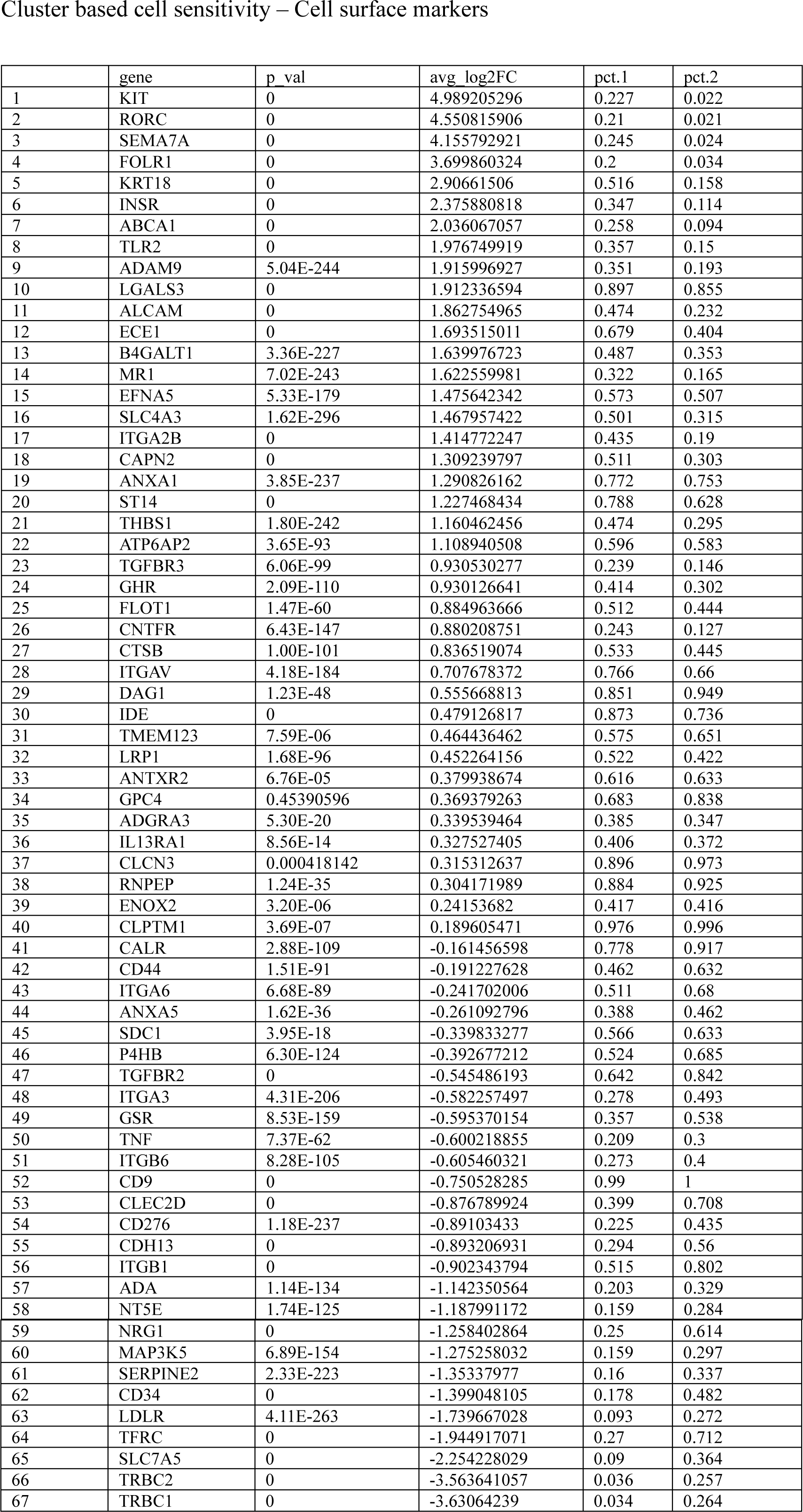
(A) List of cell surface molecules from scRNA seq which are differentially expressed in IKE responders and non-responders population

**Supplementary Figure 2.**
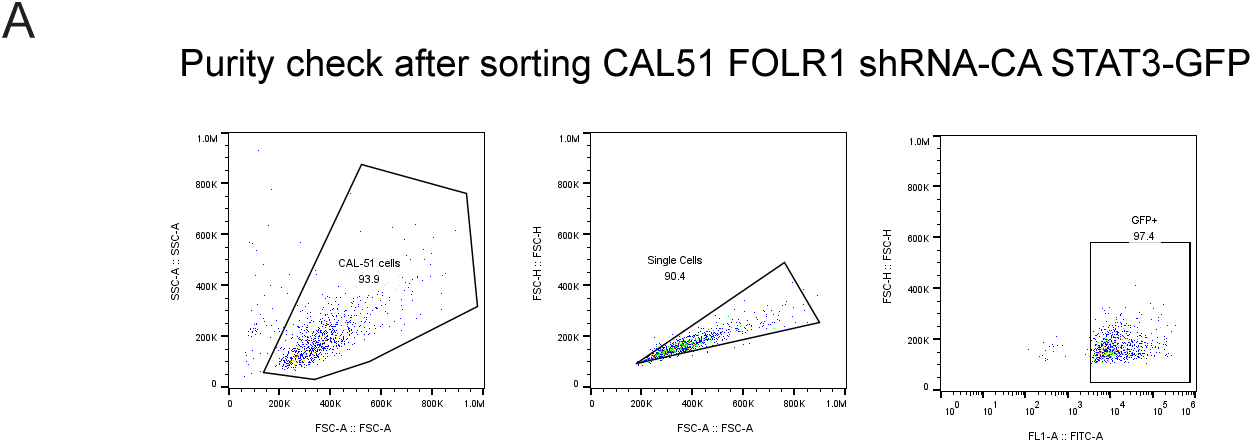
(A) Validation of constitutively active STAT3 in FOLR1 shRNA cells was performed by measuring GFP reporter by flow cytometry in GFP sorted cells.

**Supplementary Figure 3.**
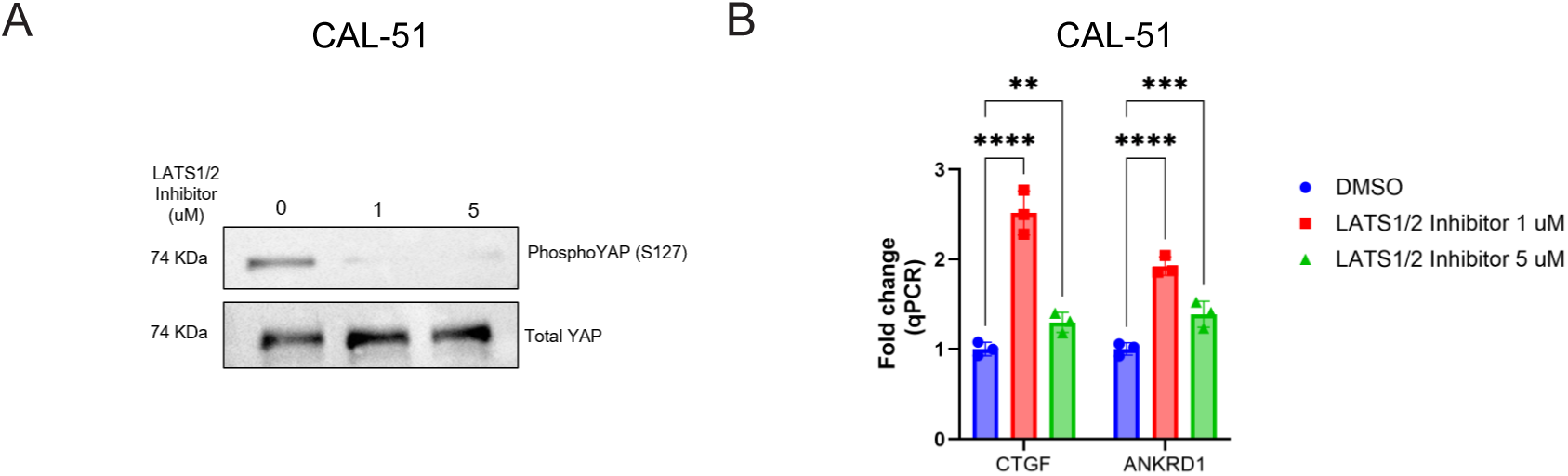
(A) Phosphorylated YAP (S127) was measured by immunoblotting in CAL-51 cells treated with DMSO or LATS1/2 inhibitor (TRULI) post 1 hr. Total YAP was used as a loading control. (B) Effect of LATS1/2 inhibitor (TRULI) on YAP target gene CTGF and ANKRD1 measured in CAL-51 cells by qPCR post 1hr treatment.

**Supplementary Figure 4.**
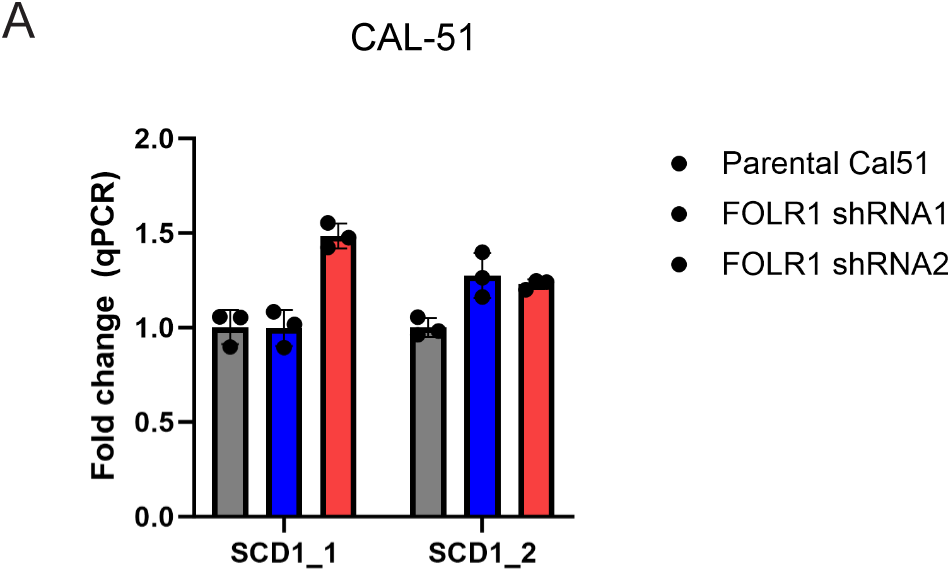
(A) SCD1 mRNA expression compared between parental CAL-51 and FOLR1 silenced cells (FOLR1 shRNA1 and FOLR1 shRNA2) by qPCR (n=3 replicates).

